# Effective generation of tumor-infiltrating lymphocyte products from metastatic non-small-cell lung cancer (NSCLC) lesions irrespective of location and previous treatments

**DOI:** 10.1101/2022.01.28.478173

**Authors:** Suzanne M. Castenmiller, Rosa de Groot, Aurélie Guislain, Kim Monkhorst, Koen J. Hartemink, Alexander A.F.A. Veenhof, Egbert F. Smit, John B.A.G. Haanen, Monika C. Wolkers

## Abstract

Non-small cell lung cancer (NSCLC) is the leading cause of cancer-related mortality worldwide. Because current treatment regimens show limited success rates, alternative therapeutic approaches are needed. We recently showed that treatment-naive, stage I/II primary NSCLC tumors contain a high percentage of tumor-reactive T cells, and that these tumor-reactive T cells can be effectively expanded and used for the generation of autologous TIL therapy. Whether these promising findings also hold true for metastatic lesions is unknown yet critical for the translation into the clinic. We here studied the lymphocytic composition and the feasibility to generate tumor-responsive TIL products from metastatic NSCLC lesions that were derived from different locations and from patients who had previously received different treatment regimens, or who were treatment naïve. The overall numbers and composition of lymphocyte infiltrates from this various metastatic lesions was by and large comparable to that of early-stage primary NSCLC tumors. With the clinically approved TIL expansion protocol, we effectively expanded TILs from metastatic NSCLC lesions to numbers that were compatible with TIL transfusion, irrespective of the location of the metastasis and of the previous treatment. Importantly, 16 of 21 (76%) tested TIL products displayed anti-tumoral activity, as determined by their production of the key pro-inflammatory cytokines IFN-γ, TNF, IL-2 and of CD137 expression upon incubation with autologous tumor digests by CD8^+^ and CD4^+^ T cells. In summary, metastatic NSCLC lesions constitute a viable source for the generation of tumor-reactive TIL products for therapeutic purposes irrespective of their location and the pretreatment regimens.

## Introduction

Lung cancer is the most common cause of cancer-related mortality world-wide^1^. Non-smallcell lung cancer (NSCLC) accounts for 85% of the patients diagnosed with lung cancer^2^. Several new treatment strategies such as targeted therapy^3,4^, immunotherapy ^5–7^ or a combination of chemo- and immunotherapy have been developed^8–11^. Nevertheless, the 5-year survival rate for NSCLC patients is still dismal^12^. Therefore, alternative treatment options for late-stage NSCLC patients are required.

Due to their high somatic mutational rate, NSCLC tumors are considered immunogenic^13^. Indeed, we and others reported a high percentage of T cell infiltrates in NSCLC lesions from treatment-naïve patients^14–16^, and evidence that these tumor-infiltrating T cells (TILs) are tumor-specific and cytotoxic is accumulating^17,18^. Adoptive TIL therapy for NSCLC patients could thus represent a viable alternative to current treatment regiments. TIL therapy is based on 4-6 weeks *in vitro* expansion of autologous TILs derived from tumor lesions that are then re-infused into pre-conditioned patients^19^. In stage III/IV melanoma patients, adoptive TIL therapy achieved an astounding overall response rate of over 50% and a complete response rate of 20%^20–23^. The similarly high immunogenicity of NSCLC implies that also NSCLC patients could also benefit from TIL therapy.

We previously reported that tumor-reactive TIL products can be efficiently generated from treatment-naïve early-stage primary NSCLC tumor lesions^14^. Importantly, more than 70% of the TIL products contained tumor-reactive T cells that produced at least one of the key pro-inflammatory cytokines Interferon gamma (IFN-γ), Tumor Necrosis Factor (TNF) and/or Interleukin-2 (IL-2) when exposed to autologous tumor digest^14^. In addition, T cells showed increased expression of the co-stimulatory receptor CD137 (also referred to as 4-1BB)^14^, which is indicative of T cell receptor triggering in this co-culture system^14,24^. Intriguingly, 25% of the expanded TILs were polyfunctional and produced more than one cytokine^14^, a feature that is correlated with the presence of highly functional effector T cells^25,26^. Our study thus demonstrated that tumor-reactive TIL products can be readily generated from treatment-naive primary NSCLC tumor lesions.

Importantly, a recent phase I clinical trial for TIL therapy in late-stage αPD-1 refractory NSCLC patients showed striking therapeutic effects^27^. In 11 out of 16 patients (68.8%), tumor regression was observed within one month after treatment^27^. Two patients (12.5%) had a complete response that was ongoing for at least 1,5 years^27^. These exciting results thus highlight the potential of TIL therapy for NSCLC patients. However, to effectively implement TIL therapy for NSCLC patients, it is paramount to define which patient groups are likely to benefit from this treatment. Patients receive are given different treatments (pre-treatment regimen), such as (immuno)chemotherapy, immune checkpoint inhibitors or small molecule inhibitors. Similarly, the location of metastatic tumor lesions^28^ may influence the potential of TILs to expand and to respond to autologous tumors. Collecting such data is thus critical for the clinical implementation of TIL therapy for NSCLC patients.

In this study, we provide an in-depth analysis of the lymphocyte composition of metastatic lesions from 27 late-stage NSCLC patients originating from different metastatic locations, and from patients who received various pre-treatment regimens, or were treatment naïve. We show that TILs can be effectively expanded from all metastatic tumor lesions. Importantly, most TIL products contained tumor-reactive T cells, as defined by cytokine production in response to autologous tumor digests. In conclusion, our study demonstrates that tumor-reactive TIL products can be generated for therapeutic purposes from late-stage NSCLC lesions, irrespective of the pre-treatment regimen, and the location of the metastatic tumor lesion.

## Materials and Methods

### Patient cohort and study design

Between October 2017 and April 2020, 27 late-stage NSCLC patients were included in this study. Characteristics of the patients, origin of metastasis, pre-treatment regimens and time between last treatment and surgery are depicted in Table 1. The cohort consisted of 13 male and 14 female patients between the age of 39 and 87 years (average 60.2 years) with clinical stage III (n=1) and stage IV (n=26) patients according to the TNM7 staging system for NSCLC. 20 patients (74%) had a history of smoking. The study protocol was designed to define the phenotype of TILs isolated from late-stage NSCLC (n=25), to explore the capacity to expand TILs (n=27), and to measure the presence of tumor-reactive T cells within TIL products (n=22). The study was performed according to the Declaration of Helsinki (seventh revision, 2013), with consent of the Institutional Review Board of the Netherlands Cancer Institute - Antoni van Leeuwenhoek Hospital (NKI-AvL), Amsterdam, the Netherlands. Tumor tissue was obtained directly after surgery, transported at room temperature in RPMI medium 1640 (Gibco) containing 50 μg/ml gentamycin (Sigma-Aldrich), 12,5 μg/ml Fungizone (Amphotericin B, Gibco) and 20% fetal bovine serum (FBS, Bodego), weighed and processed within four hours. Tumor stage and differentiation, weight of the obtained tissues and total tumor weight for each patient are shown in Table 2. Since surgery was not always performed for diagnostic purposes, not all tumor characteristics were determined for each patient. Tumor size ranged from T1a-T4 according to the TNM7 system, and specified as adenocarcinoma (AC, n = 23), squamous carcinoma (SCC, n = 2) and as NSCLC not otherwise specified (NOS, n = 2). On average, 1610 mg tumor tissue was obtained for this study, ranging from 120 - 8810 mg.

**Table 1:** Patient characteristics.

| Patient | Gender (M/F) | Age at surgery (years) | Smoking history (cigarettes per day) | Clinical stage (TNM7*) | Tumor lesion site | Pre-treatment regimen | Last treatment (months before surgery) |
| --- | --- | --- | --- | --- | --- | --- | --- |
| 1 | M | 64 | Unknown | IVA | Lung | Nivolumab + Ipilimumab (immunotherapy) | 6 |
| 2 | F | 64 | No (never) | IVA | Lung | cis/pem + carbo/pem (chemotherapy) | 0 |
| 3 | M | 45 | Yes (unknown) | IIIA | Lung/Lymph node | Treatment naïve |  |
| 4 | M | 55 | Yes (16/d) | IV | Lymph node | Treatment naïve |  |
| 5 | F | 48 | Yes (20/d) | IV | Lung | cis/pem (chemotherapy) | unknown |
| 6 | F | 50 | Yes (20/d) | IV | Lymph node | Nivolumab (immunotherapy) | 4 |
| 7 | M | 58 | Yes (6/d) | IV | Lung | Pembrolizumab (immunotherapy) | 1 |
| 8 | F | 57 | No (never) | IV | Lung | Pembrolizumab (immunotherapy) | 1 |
| 9 | M | 55 | No (never) | IV | Lung | Erlotinib/osimertinib (EGFR inhibitor) | 0 |
| 10 | F | 39 | Yes (25/d) | IV | Adrenal gland | Pembrolizumab (immunotherapy) | 0 |
| 11 | F | 60 | Yes (13/d) | IV | Subcutaneous tissue - hip | Treatment naïve |  |
| 12 | F | 69 | No (never) | IV | Liver | EGFR inhibitor | 0 |
| 13 | M | 53 | No (never) | IV | Adrenal gland | Alectinib (ALK inhibitor) | 15 |
| 14 | F | 70 | Yes (20/d) | IV | Adrenal gland | carbo/pem (chemotherapy) | 17 |
| 15 | M | 66 | Yes (unknown) | IV | Subcutaneous tissue - skin | cis/gem + Nivolumab (chemo-immuno) | 1 (immuno) |
| 16 | M | 87 | Yes (8/d) | IV | Adrenal gland | Crizotinib (ALK inhibitor) | 0 |
| 17 | F | 73 | No (never) | IV | Adrenal gland | Osimertinib (EGFR inhibitor) | continued |
| 18 | M | 60 | Yes (30/d) | IV | Spleen | CRT (chemo) + Nivolumab (immunotherapy) | 0 (immuno) |
| 19 | F | 47 | Yes (5/d) | IV | Adrenal gland | Nivolumab + anti-LAG3 (immunotherapy) | 0 |
| 20 | M | 55 | Yes (4/d) | IV | Lung | carbo/pem + Pembrolizumab (chemo-immuno) | 1 (immuno) |
| 21 | F | 40 | Yes (25/d) | IV | Adrenal gland | cis/gem + Pembrolizumab (chemo-immuno) | 0 (immuno) |
| 22 | M | 58 | Yes (4/d) | IV | Lung | cis/eto/carbo + Nivolumab (chemo-immuno) | 0 (immuno) |
| 23 | F | 66 | Yes (19/d) | IV | Lung | Pembrolizumab (immunotherapy) | 1 |
| 24 | M | 77 | Yes (10/d) | IV | Lung | Treatment naïve |  |
| 25 | M | 73 | Yes (10/d) | IV | Adrenal gland | carbo/pem (chemotherapy) | 14 |
| 26 | F | 65 | Yes (25/d) | IV | Lung | Pembrolizumab (immunotherapy) | 1 |
| 27 | F | 72 | Yes (5/d) | IV | Lung | cis/pem (chemotherapy) | unknown |

\*American Joint committee on Cancer: Lung Cancer Staging, 7th edition, 2009
M = Male; F = Female; d = day

**Table 2:** Tumor characteristics.

| **Donor** | **Tumor differentiation** | **Pathology stage (TNM7\*)** | **PDL-1 status (determined at PA)** | **Biopsy weight (mg)** | **Tumor size (cm3)** |
| --- | --- | --- | --- | --- | --- |
| **1** | AC | ypT2bN0PL0/cT4N3M1a | N.D. | 2140 | 4.7 |
| **2** | AC | ypT2aN2PL0/dT4N2M1a | 80% | 670 | 3.2 |
| **3** | AC | pT1bN2PL0 | 100% | 1500 | 1.5 |
| **4** | AC | N.D. | 70% | 1408 | 2.0 |
| **5** | NOS | cT3N1M1c/ypTON0 | N.D. | 320 | 1.5 |
| **6** | AC | cT3N3M1b | 1% | 1000 | 3.5 |
| **7** | AC | N.D. | 90% | 230 | 2.4 |
| **8** | AC | ypT1aN0PL0 | 5% | 800 | 1.0 |
| **9** | AC | ypT1cN0M1aPL0 | <1% | 500 | 2.5 |
| **10** | AC | cT3N2M1 | <1% | 600 | 2.6 |
| **11** | AC | N.D. | <1% | 1200 | 1.2 |
| **12** | AC | N.D. | <1% | 4000 | 4.6 |
| **13** | AC | N.D. | 100% | 250 | 3.0 |
| **14** | NOS | cT1cN2M1c | 70% | 1810 | 7.5 |
| **15** | SSC | cTxN3M1b | <1% | 8810 | 4.5 |
| **16** | AC | N.D. | 30% | unknown | 7.5 |
| **17** | AC | pT2N2PL1M0 | 1% | unknown | 2.8 |
| **18** | AC | N.D. | 80% | 8590 | 7.1 |
| **19** | AC | N.D. | 2% | 580 | 3.5 |
| **20** | AC | ypTON1 | 70% | 590 | unknown |
| **21** | AC | cT3N2M1 | <1% | unknown | 1.7 |
| **22** | SSC | pT2bN0PL1 | 50% | unknown | 4.2 |
| **23** | AC | pT1bPL0 | 90% | 150 | 1.9 |
| **24** | AC | pT4N2PL1 | N.D. | 600 | 3.2 |
| **25** | AC | cT1cN0M2 | <1% | 750 | 3.7 |
| **26** | AC | N.D. | <1% | 120 | 1.4 |
| **27** | AC | ypT3PL1 | <1% | 420 | 1.2 |

\*American Joint committee on Cancer: Lung Cancer Staging, 7th edition, 2009
AC = adenocarcinoma; SCC = squamous carcinoma; NOS = NSCLC not otherwise specified; PA = pathology
N.D. = not determined

### Tissue digestion

Tissue digestion was performed as previously reported^14^. Briefly, freshly isolated tumorous tissue was finely chopped and incubated for 45 minutes on a rotor at 37°C in RPMI medium 1640 containing 30 IU/ml collagenase IV (Worthington), 12,5 μg/ml DNAse (Roche), and 1% FBS. The digest was pelleted at 360 g for 10 min and resuspended in FACS buffer (PBS containing 2% FBS and 2 mM EDTA). The digest was filtered through a tea mesh and then over a 100 μm filter. After red blood cell lysis for 15 minutes at 4°C with 155 mM NH4Cl, 10 mM KHCO3, 0.1 mM EDTA (pH 7.4), live and dead cells were manually counted with trypan blue solution (Sigma) on the haemocytometer. 1-2×10^6^ live cells were used for flow cytometry analysis, and 1-3×10^6^ live cells were used for TIL cultures. The remaining digest was cryopreserved until further use.

### T cell expansion

For TIL expansion, 2-3 wells containing 0.5-1 x10^6^ live cells from the tissue digest were cultured for 10-13 days in 24 wells plates in 20/80 T-cell mixed media (Miltenyi) containing 5% human serum (HS) (Sanquin), 5% FBS, 50 μg/ml gentamycin, 1.25 μg/ml fungizone, and 6000 IU human recombinant (hr) IL-2 (Proleukin, Novartis) (pre-Rapid Expansion Phase; pre-REP) at 37°C and 5% CO_2_. Medium was refreshed on day 7, 9 and 11 of the preREP phase. When a monolayer of cells was observed in the entire well, cells were split in 2 wells. When more than 30% of the cells stopped dividing (determined as rounding of cells), cells were harvested, counted and prepared for an additional culture period of 10–13 days (REP). 2×10^5^ live cells/well (1-3 wells/patient) were co-cultured with 5-10×10^6^ irradiated PBMCs pooled from 15 healthy blood donors in a 24-wells plate, 30 ng/ml anti-CD3 antibody (OKT-3) (Miltenyi Biotec) and 3000 IU/ml hrIL-2. Typically, cells were passaged when a monolayer of cells was observed in the wells during the REP phase (generally every other day) and harvested, washed and counted on day 10–13 after REP, based on visual assessment of the T cell proliferation state as mentioned above. Cells were either used immediately or were cryo-preserved in RPMI medium 1640 containing 10% Dimethyl Sulfoxide (DMSO) (Corning) and 40% FBS until further use.

### T cell activation

When enough material was available, fresh or defrosted REP TILs were counted and prestained in FACS buffer with anti-CD4 BUV496 and anti-CD8 BUV805 (BD Biosciences) for 30 minutes at 4°C. Cells were washed twice, i.e. once with FACS buffer, and once with 20/80 T-cell mixed media (Miltenyi). 1×10^5^ live expanded TILs were co-cultured with 2×10^5^ live tumor digest cells for 6 hours at 37°C. Alternatively, expanded TILs were stimulated with 10 ng/ml PMA and 1 μg/ml Ionomycin (SigmaAldrich), or cultured with T-cell mixed media alone. After 1 hour of co-culture, 1x Brefeldin A (Invitrogen) and 1x Monensin (Invitrogen) were added.

### Flow cytometry

For ex vivo analysis, tumor digests were washed twice with FACS buffer and stained in FACS-buffer for 30 minutes at 4°C with the following antibodies: anti-CD3 PerCp-Cy5.5, anti-CD279 FITC, anti-CD103 FITC, anti-CD56 BV605, anti-CD27 BV510, anti-CD127 BV421, anti-CD279 BV421, anti-CD39 PE-Cy7, anti-CD103 PE-Cy7 and anti-CD25 PE (Biolegend), and with anti-CD8 BUV805, anti-CD45RA BUV737, anti-CD4 BUV496, and anti-CD69 BUV395 (BD Biosciences). Live/dead fixable near IR APC-Cy7 (Invitrogen) was included for dead cell exclusion. Cells were washed twice with FACS buffer and fixed for 30 minutes with the Perm/Fix Foxp3 staining kit (Invitrogen) according to the manufacturer’s protocol. Cells were stained with anti-Foxp3 Alexa647, anti-CD137 Alexa647 or anti-CD137 PE-Cy7 (Biolegend) for 30 minutes at 4°C. Cells were resuspended in FACS buffer and passed through a 70 μm single cell filter prior to flow cytometry analysis (Symphony A5, BD Biosciences). Expanded TILs were washed twice with FACS buffer and then pre-stained with anti-CD4 BUV496 and anti-CD8 BUV805 as described above before activation. Anti-CD107 Alexa700 (BD Biosciences) was included during the coculture with tumor cells. After T cell activation, cells were washed twice with FACS buffer and stained with anti-CD3 PerCp-Cy5.5 and anti-CD279 BV421 (Biolegend) and Live/dead fixable near IR APC-Cy7 in FACS buffer for 30 minutes at 4°C. After two washes with FACS buffer, cells were fixed with Perm/Fix Foxp3 staining kit (Invitrogen) and then stained with anti-CD137 Alexa647 (Invitrogen), anti-CD154 BV510, anti-IFN-γ PE, anti-TNF-α BV785 and anti-IL-2 FITC (Biolegend) in PermWash buffer according to the manufacturer’s protocol. Cells were washed with FACS buffer and passed through a 70 μM single cell filter prior to acquisition with the Symphony A5 flow cytometer (BD). Flow cytometry settings were defined for each patient with single staining for each antibody. To each analysis performed, a standardized sample of PBMCs pooled from four healthy donors that was cryo-preserved prior to the start of the study was included. Data analysis was performed with FlowJo Star 10.7.1.

### Statistical analysis

Statistical analysis was performed with GraphPad Prism 8.0.2. Compiled data are shown as paired data points for each patient, or as single data points with box and whiskers showing maximum, 75th percentile, median, 25th percentile and minimum, unless otherwise stated in the legend. The overall significance was calculated with unpaired parametric t test with a twotailed p-value, with ordinary 2-way ANOVA test, or with paired parametric t test with a two-tailed p-value, variance was calculated as standard deviation (S.D.). The p-value cut-offs were set on * = p<0.05, ** = p<0.01, *** = p<0.001, **** = p<0.0001.

## Results

### Late-stage metastatic NSCLC tumor lesions contain high numbers of T cell infiltrates

To determine the rate of viable cells and specifically of T cell infiltrates within metastatic NSCLC tumor tissues, we processed tumor tissues from 27 late-stage NSCLC patients to single cell suspensions with the clinically approved digestion protocol and compared that to our previous study from early-stage primary NSCLC tumors^14^. We collected on average 41.2 x 10^6^ (range 1.5-206 x 10^6^) viable immune cells per gram tissue, which is comparable to the yield of viable immune cells we achieved from early-stage primary tumor lesions^14^ (figure 1(a)). Flow cytometric analysis revealed that the numbers of lymphocytes per gram tissue in metastatic lesions reached on average 17.0 x 10^6^ cells (range 0.6-82.1×10^6^ cells) (figure S1, S2), and the majority thereof (71.4% ± 21.4%) consisted of CD3^+^ T cells, with on average 15.5 x 10^6^ cells (range 0.2-70.0 x 10^6^) per gram tissue (figure 1(a)). The average yield in absolute numbers and the percentage of CD3^+^ T cell infiltrates was thus similar to that of early-stage tumor lesions^14^ (figure 1(a)), yet displayed a broader range. We therefore assessed if the pre-treatment regimens or the location of the metastatic tumor lesion influenced the T cell content. Specifically, the cohort contained lesions from treatment-naïve (n=4) patients, and from patients that had received different (combinatory) pre-treatment regimens, i.e. immunotherapy (n=8), chemotherapy (n=5), a combination of those two (n=5), or small molecule inhibitors, like anaplastic lymphoma kinase (ALK) inhibitors (n=2) and epidermal growth factor receptor (EGFR) inhibitors (n=3) (Table 1). The metastatic tumor lesions were obtained from different tissues, i.e. lung (n=12), lymph node (n=3), adrenal gland (n=8), liver (n=1), spleen (n=1), and subcutaneous tissue (n=2) (Table 1). Neither metastatic location nor pre-treatment regimen showed clear differences in the yield of viable cells and/or of CD3^+^ T cell infiltrates (figure 1(b, c)). Thus, CD3^+^ T cells can be readily obtained from metastatic NSCLC tumor lesions independently of treatment regimens the patient previously received and the location of metastases.

**Figure 1.**
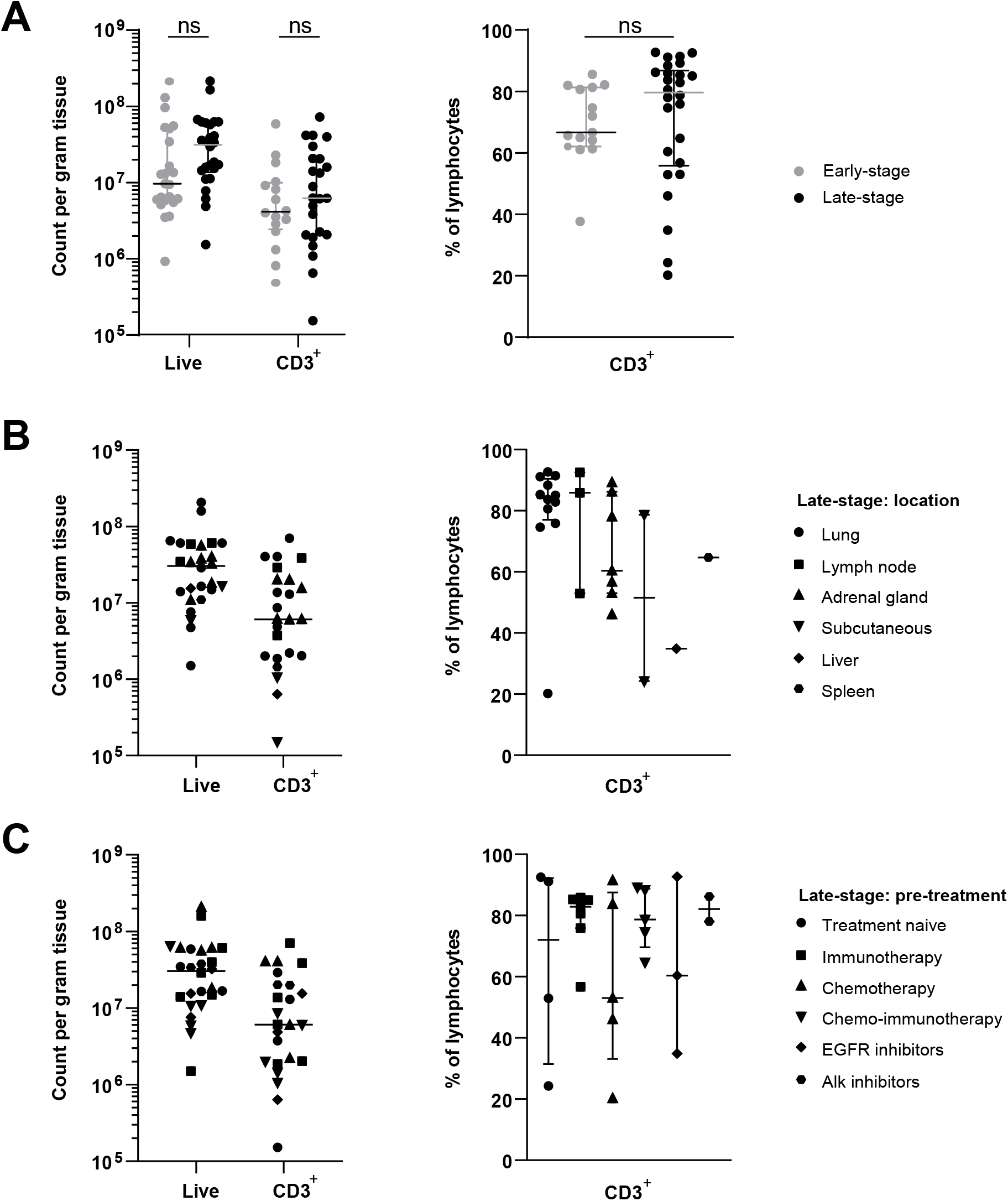
Effective isolation of lymphocytes from late-stage NSCLC lesions. Single cell suspensions were obtained by enzymatic digestion of metastatic lesions from late-stage NSCLC patients. Life cell count was assessed with tryphan blue staining, and lymphocyte and CD3^+^ percentage was determined by flow cytometry. (A-C) Left panels: count of live and of CD3^+^ T cells per gram tissue. Right panels: percentage of CD3^+^ T cells within the lymphocyte population. (A) Composition of the yield of live and CD3^+^ T cells in tumor lesions from the late-stage NSCLC patients (n=27, black) compared to treatment-naïve primary tumor lesions as reported previously (n=17, grey)^14^. (B) Comparison of the yield of live and of CD3^+^T cells from different tumor lesion sites, indicated as lung (n=12), lymph node (n=3), adrenal gland (n=8), subcutaneous tissue (n=2), liver (n=1) and spleen (n=1). (C) Comparison of different pretreatment regimens, indicated as treatment-naïve (n=4), immunotherapy (n=8), chemotherapy (n=5), chemo-immunotherapy (n=5), EGFR inhibitors and ALK inhibitors(n=2). Each dot represents one patient. All graphs show median, A and B,C right panel also depicts 25th and 75th pct interval. Significance was calculated with unpaired student’s t test. Not significant (ns): p ≥ 0.05.

### Conserved lymphocyte infiltration profile in late-stage metastatic NSCLC lesions

We next questioned how the lymphocyte composition in metastatic NSCLC tumor lesions compared to that of early-stage NSCLC primary tumor lesions (gating strategy: figure 2(a))^14^. We therefore measured the percentage of T cell populations, B cells, NK cells and NKT cells. Overall, the distribution of lymphocyte populations in late-stage metastatic tumor lesions was comparable to that in primary early-stage tumor lesions (figure 2(b)). CD3^+^ T cells were most prevalent, with 31.8% ± 16.6% CD8^+^ T cells, 25.1% ± 8.1% conventional CD4^+^ T cells, and 7.6% ± 5.0% regulatory T cells (Tregs) (figure 2(b)). The percentage of CD8^+^ T cells and Tregs showed a broader distribution in percentages in metastatic lesions when compared to T cell infiltrates in early-stage primary tumor lesions, yet no significant differences were observed (figure 2(b)). Only the percentage of conventional CD4^+^ T cell infiltrates was significantly lower in late-stage NSCLC metastatic lesions when compared to early-stage NSCLC lesions (figure 2(b); p-value: 0.008). Even though trends for various lymphocyte distribution were observed, neither different pre-treatment regiments nor different locations of resected metastases showed overt differences in the distribution of specific lymphocyte subsets (figure 2(c)).

**Figure 2.**
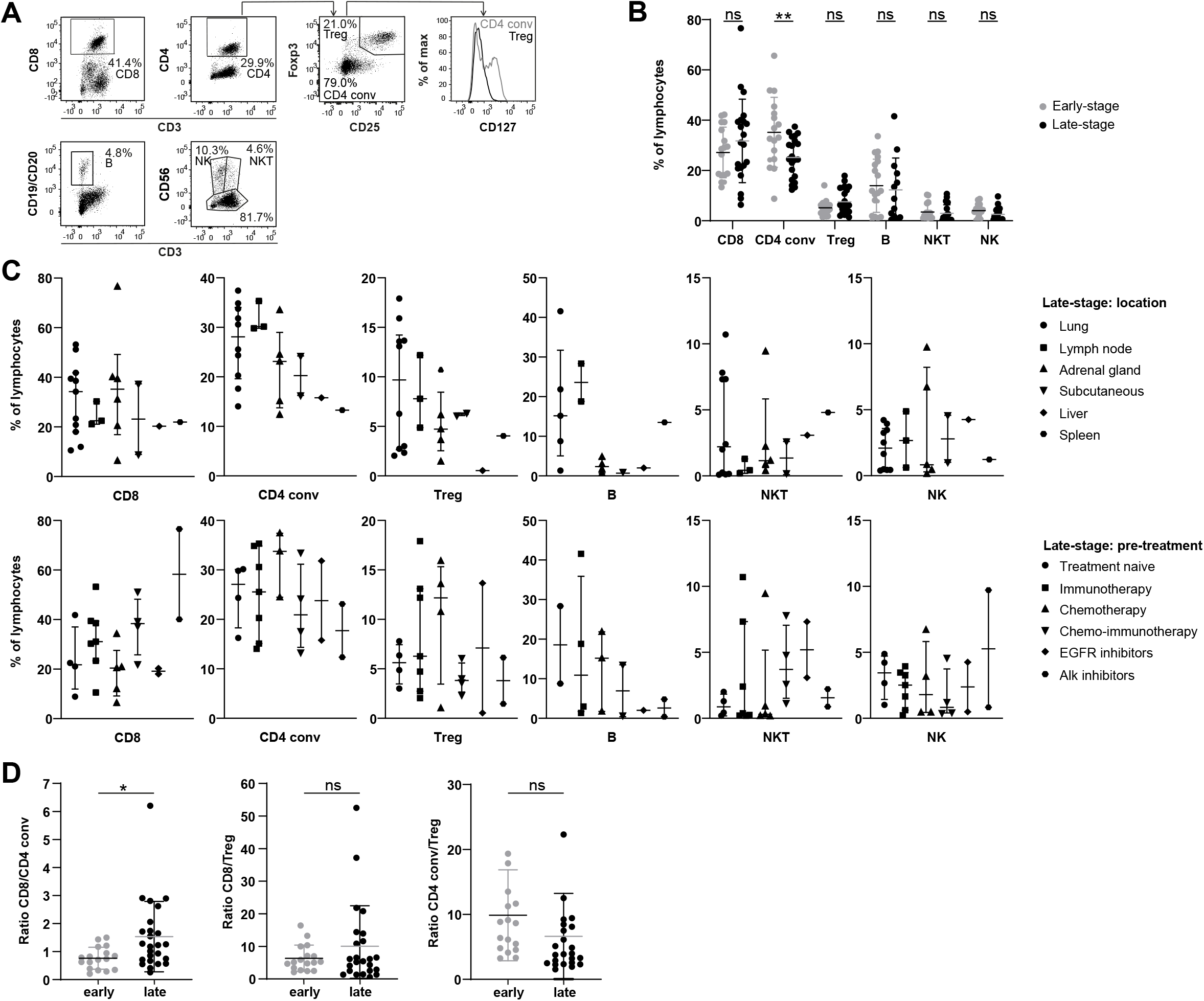
Lymphocyte composition in late-stage and early-stage tumor lesions. (A) Gating strategy for defining different lymphocyte populations (patient 13) as reported in (B-D), indicated as CD8^+^ T cells, CD4^+^ T cells, regulatory T cells, conventional CD4^+^ T cells, B cells, NK cells and NKT cells. (B) Lymphocyte composition from late-stage metastatic lesions (black, n=25) compared to early-stage primary tumor lesions (grey, n=17)^14^. (C) Percentage of indicated lymphocyte populations between patients with different tumor lesion site (top) and different pre-treatment regimens (bottom). (D) Ratio of CD8^+^ T cells over conventional CD4^+^ T cells (left panel) and regulatory CD4^+^T cells (middle panel), and of conventional CD4^+^ T cells over regulatory T cells (right panel) in late-stage NSCLC lesions (black, n=25) and early-stage primary NSCLC tumors (grey, n=17)^14^. Each dot represents one patient. All graphs show median with 25th and 75th pct interval. Significance was calculated with unpaired student’s t test.* = p<0.05, ** = p<0.01. Not significant (ns): p ≥ 0.05.

We also determined the ratio between the different T cell subsets (figure 2(d)). Whereas the ratio between CD8^+^ and regulatory T cells, and between conventional CD4^+^ and regulatory T cells did not differ between late-stage metastatic and early-stage NSCLC primary tumor lesions, we found a shift in the ratio between CD8^+^ over conventional CD4^+^ T cells in late-stage tumor lesions compared to early-stage lesions, with an average ratio of 1.54 (late-stage) compared to 0.8 (early-stage) (figure 2(d); p-value: 0.02). In conclusion, we observed a similar distribution of lymphocyte infiltrates between early-stage and late-stage tumor lesions, except for a reduction of the ratio of conventional CD4^+^ infiltrates in late-stage NSCLC tumor lesions.

### T cells from metastatic lesions display a high memory profile and high expression levels of tissue residency markers

The T cell memory profile is an important indicator for T cell function. In fact, in the early-stage NSCLC cohort, we found high percentages of effector memory T cells^14^. We therefore investigated the memory profile of T cells in metastatic lesions. To this end, we measured the expression of CD27 and CD45RA to distinguish CD27^+^CD45RA^+^ naïve (Tnaive), CD27^+^CD45RA^-^ central memory (Tcm), CD27^-^CD45RA^-^ effector memory (Tem) and CD27^-^CD45RA^+^ terminally differentiated effector memory (Temra) T cells (figure 3(a,b)). In particular the percentages of CD8^+^ Tem cells were substantially reduced in late-stage TILs when compared to the early-stage NSCLC TILs (p-value: 0.0003), concomitant with a slight but non-significant increase of percentages of CD8^+^ Tcm T cells (figure 3(a)). We observed a similar yet non-significant shift towards Tcm away from Tem in the CD4^+^ T cell compartment (figure 3(b); p-value: 0.09). The percentages of Tnaive or Temra cells in the CD8^+^ and the conventional CD4^+^ T cell compartments remained stable (figure 3(a,b)).

**Figure 3.**
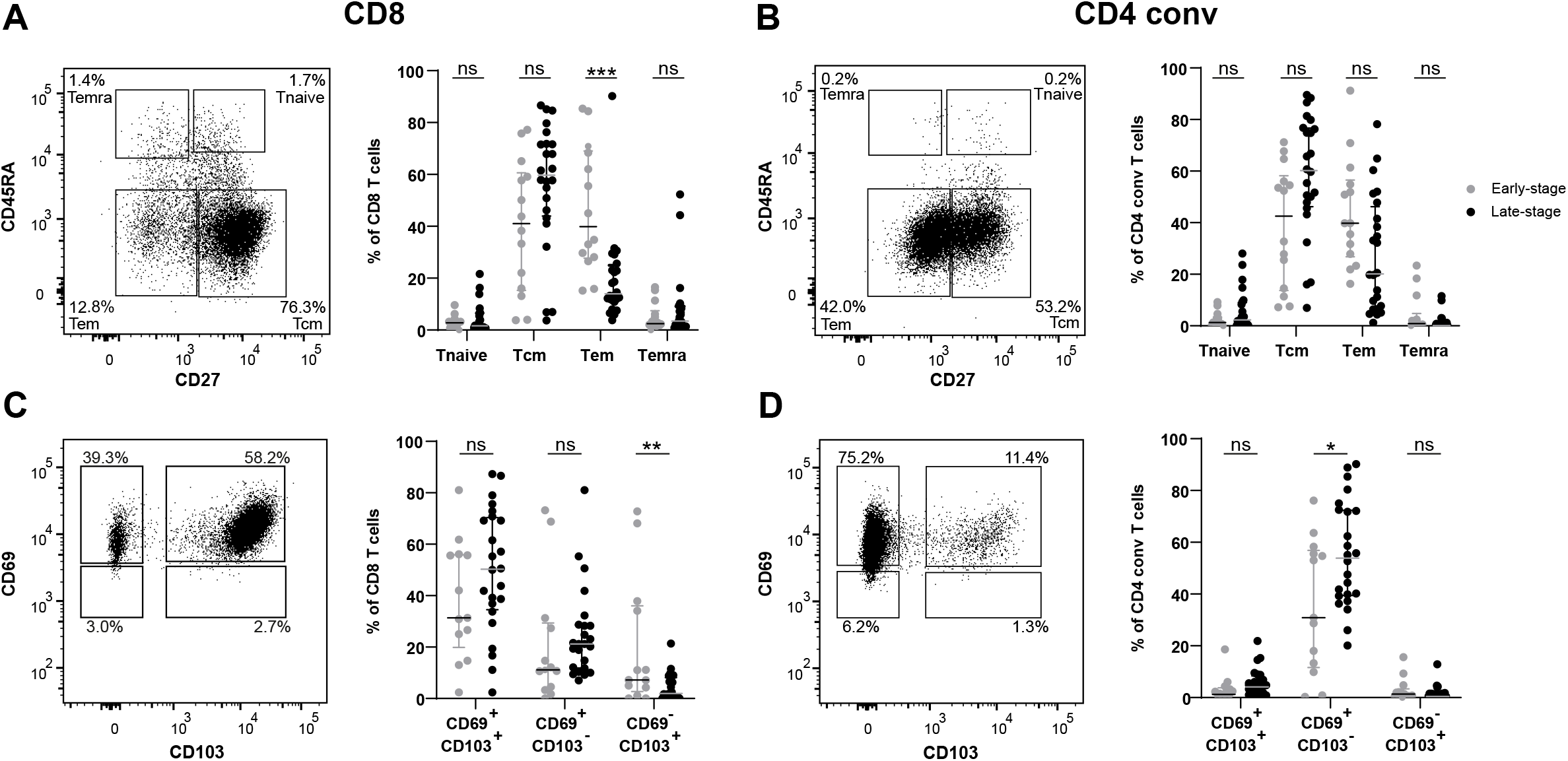
Ex vivo differentiation status and tissue residency in metastatic tumor lesions compared to primary early-stage tumor lesions. (A, B) T cell differentiation status of tumorinfiltrating T cells from late-stage (black, n=24) and early-stage (grey, n=14)^14^ tumor lesions, as defined by their expression of CD27 and CD45RA. Left: representative dot plot (patient 13). Right: The percentage of Tnaive (CD27^+^CD45RA^+^), Tcm (CD27^+^CD45RA^-^), Tem (CD27^-^CD45RA^-^) and Temra (CD27^-^CD45RA^+^) for (A) CD8^+^ T cells and (B) conventional CD4^+^ T cells. (C, D) Comparison of expression of tissue-residency markers CD69 and CD103 on TILs from late-stage (black, n=24) and early-stage (grey, n=13)^14^. Left: representative dot plot (patient 13). Right: The percentage of CD69^+^CD103^+^, CD69^+^CD103^-^ and CD69^-^CD103^+^ for (C) CD8^+^ T cells and (D) conventional CD4^+^ T cells. Each dot represents one patient. All graphs show median and 25^th^ and 75^th^ pct interval. Significance was calculated with unpaired student’s t test. * = p<0.05, ** = p<0.01, *** = p<0.001. Not significant (ns): p ≥ 0.05.

We also measured the expression levels of the two tissue residency markers CD103 and CD69 (figure 3(c,d)). Tissue residency markers can be used as a clinical parameter for patients with lung tumor lesions^29^. We found high percentages of CD69^+^CD103^+^ (49.9% ± 23.9%) and CD69^+^CD103^-^ (25.1% ± 17.3%) CD8^+^ T cells and low numbers of CD69^-^CD103^+^ (4.3% ± 5.3%) CD8^+^ T cells (figure 3(c)). In line with literature^14,30^, most conventional CD4^+^ T cells lack the expression of CD103 as tissue residency marker, and contain rather high percentages of CD69 expressing cells, a feature that is also observed in late-stage tumor lesions (figure 3(d)). We found the majority of conventional CD4^+^ T cells expressing CD69^+^CD103^-^ (55.3% ± 20.3%), and low number of CD69^+^CD103^+^ (6.0% ± 5.2%) and CD69^-^CD103^+^ (1.7% ± 2.7%). Intriguingly, both CD8^+^ and CD4^+^ T cells displayed higher percentages of T cells expressing tissue residency markers in late-stage tumors compared to early-stage tumor lesions (figure 3(c, d)). Thus, late-stage TILs display a decrease in the Tem compartment and an increase in tissue residency traits when compared to early-stage TILs.

### Tissue resident T cells from metastatic lesions express high levels of activation markers

To further define the fitness of the T cell infiltrates from late-stage NSCLC tumor lesions, we measured several exhaustion and activation markers with a multi-color flow cytometry panel. We first measured the expression of Programmed Death receptor 1 (PD-1), a marker that correlates with the dysfunctional state of TILs^31,32^. In late-stage TILs, the PD1 levels reached 10.2% ± 14.4 % for CD8^+^ T cells (figure 4(a)) and 8.8% ± 9.4 % for conventional CD4^+^ T cells (figure 4(b)), respectively. These percentages did not significantly differ from those of early-stage primary NSCLC tumors (figure 4(a,b)).

**Figure 4.**
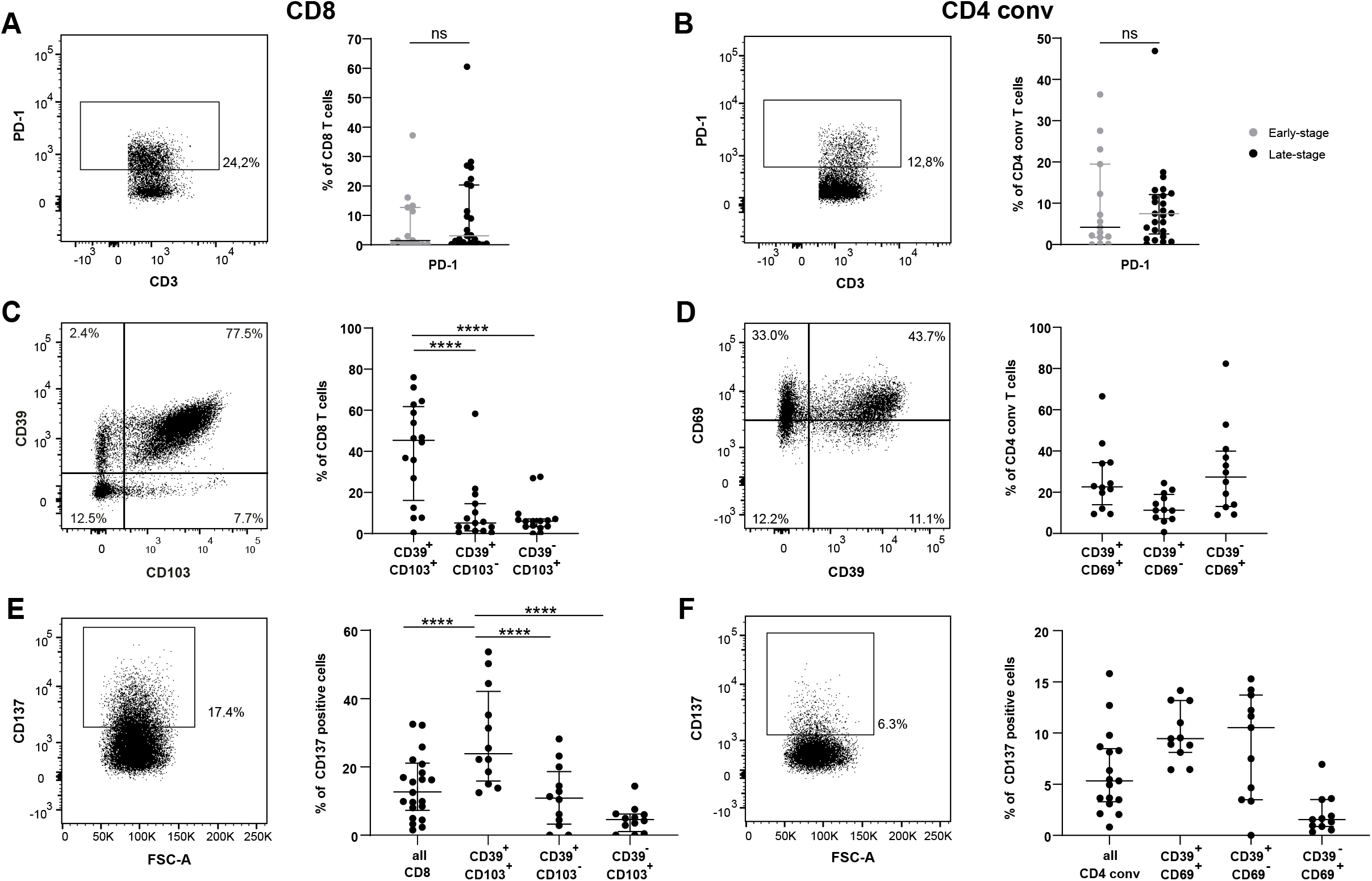
Ex vivo late-stage TILs express activation and exhaustion markers on both CD8^+^ T cells and CD4^+^ conventional T cells. (A, B) PD-1 expression between late-stage (black, n=25) and early-stage (grey, n=15)^14^ TILs. Left: Gating strategy for defining PD1 expression (patient 13). Right: percentage of PD-1 for (A) CD8^+^ T cells and (B) conventional CD4^+^ T cells. (C,D) CD39 expression in tissue-resident T cells. Left: Gating strategy for defining CD39 expression (patient 13). Right: percentage of (C) CD39 and CD103 expression for CD8^+^ T cells and (D) CD39 and CD69 expression for conventional CD4^+^ T cells. (E,F) CD137 (4-IBB) expression, calculated as percentage of the subpopulation labelled on the X-axis. Left: Gating strategy for defining CD137 expression (patient 13). Right: CD137 expression in (E) CD39 and CD103 expressing cells for CD8^+^ T cells and in (F) CD39 and CD69 expressing cells for conventional CD4^+^ T cells. Each dot represents one patient. All graphs show median and 25^th^ and 75^th^ pct interval. Significance was calculated with 2-way ANOVA test (Tukey’s Multiple Comparison). *** = p<0.001, **** = p<0.0001. If no indication, p ≥ 0.05.

Another surface protein that marks activated T cells and exhausted T cells is the ectonucleoside triphosphate diphosphohydrolase-1 CD39^33–35^. In particular when combined with the tissue residency markers CD69 and CD103, CD39 was found to identify T cells with the capacity to infiltrate human solid tumors^35–38^. We therefore measured CD39 expression in combination with CD103 expression in CD8^+^ T cells (figure 4(c)). CD39 was most prominently expressed in CD103^+^ T cells, with an average expression of 41.5% ± 24.0% for CD39^+^CD103^+^ CD8^+^ T cells compared to 11.1% ± 14.9% for CD39^+^CD103^-^ (p-value <0.0001) and 8.5% ± 8.3% for CD39^-^CD103^+^ (p-value <0.0001). Because conventional CD4^+^ T cells barely expressed CD103 (figure 3(d)), we examined the CD39 expression in combination with expression of CD69 (figure 4(d)). In the conventional CD4^+^ T cells, the CD39 expression is more evenly distributed between the different populations, with 26.7% ± 16.3% of the conventional CD4^+^ T cells being CD39^+^CD69^+^, 12.7% ± 7.0% being CD39^+^CD69^-^ and 30.4% ± 21.3% being CD39^-^CD69^+^(figure 4(d)).

To further define the activation status of the TILs, we measured the expression of the costimulatory molecule CD137 expression. CD137 expression was shown to enrich for naturally occurring tumor-reactive T cells^39^. In metastatic NSCLC lesions, we found 13.9% ± 9.1% of the tumor infiltrating CD8^+^ T cells and 6.1% ± 4.0% of the conventional CD4^+^ T cells expressed CD137 (figure 4(e,f)). Intriguingly, tumor infiltrating CD8^+^ T cells that express the tissue residency marker CD103 together with CD39 showed with 28.7% ± 14.3% a substantial increase in the percentage of CD137 expression (figure 4(e); p-value < 0.0001). We made similar observations when we used CD69 expression as a marker for tissue residency (figure S3(a)). Albeit not significant, similar trends of increased CD137 expression were detected on CD69^+^ tissue resident cells conventional CD4^+^ T cells with or without CD39 expression. Specifically, CD137 expression increased to 9.9% ± 2.7% on the CD39^+^CD69^+^ conventional CD4^+^ T cells and to 8.8% ± 5.2 of the CD39 single positive conventional CD4^+^ T cells, compared to 2.1% ± 1.9% CD69 single positive conventional CD4^+^ T cells (figure 4(f)). In conclusion, tissue resident CD8^+^ CD4^+^ T cells show increased levels of CD39 and CD137.

Overall, we measured higher percentages of Tcm cells at the expense of Tem cells in late-stage NSCLC tumor lesions compared to TILs from early-stage NSCLC tumor lesions. Tissue residency markers are increased on CD8^+^ and CD4^+^ tumor-infiltrating T cells. Furthermore, the expression of the activation markers CD39 and CD137 are increased, which all combined suggests that tumor-responsive T cells are present in metastatic NSCLC lesions.

### Effective expansion of TILs from late-stage NSCLC tumor lesions

We next determined how efficient TILs could be expanded from the metastatic lesions with the clinically approved rapid expansion protocol (REP), the current golden standard for generating TIL products for the clinic^22^. During the first 10-13 days of culturing tumor digests with human recombinant IL-2 (pre-REP), the metastatic TILs expanded on average a 40-fold (figure 5(a)). The following REP-phase of 10-13 days in the presence of CD3 stimulation and IL-2 resulted in an additional 100-fold expansion of TILs (figure 5(a)), and a total expansion rate of on average 3500-fold (figure 5(a)). Late-stage TILs expanded slightly more during the pre-REP phase than early-stage TILs, and less during the REP phase (figure 5(a)). Yet, the total expansion rate from these two cohorts was comparable (figure 5(a))^14^. Furthermore, the expansion rate did not substantially differ for TILs originating from different metastatic locations (figure 5(b)) or from patients that underwent different pre-treatments (figure 5(c)). Also the amount of total viable cells per gram tissue we obtained after expansion is comparable with those we reported for early-stage tumors (figure 5(d))^14^. Notably, when we used the tumor size (Table 2) to extrapolate the tumor mass and to estimate the expansion rate from the total tumor lesion, all 27 TIL products would have resulted in > 1 x 10^9^ cells (figure 5(d)), which is compatible with TIL infusion into patients. Thus, TILs can be efficiently expanded from late-stage metastatic NSCLC lesions, and this expansion does not depend on the location of the metastatic tumor and/or the previous treatments given to the patient.

**Figure 5.**
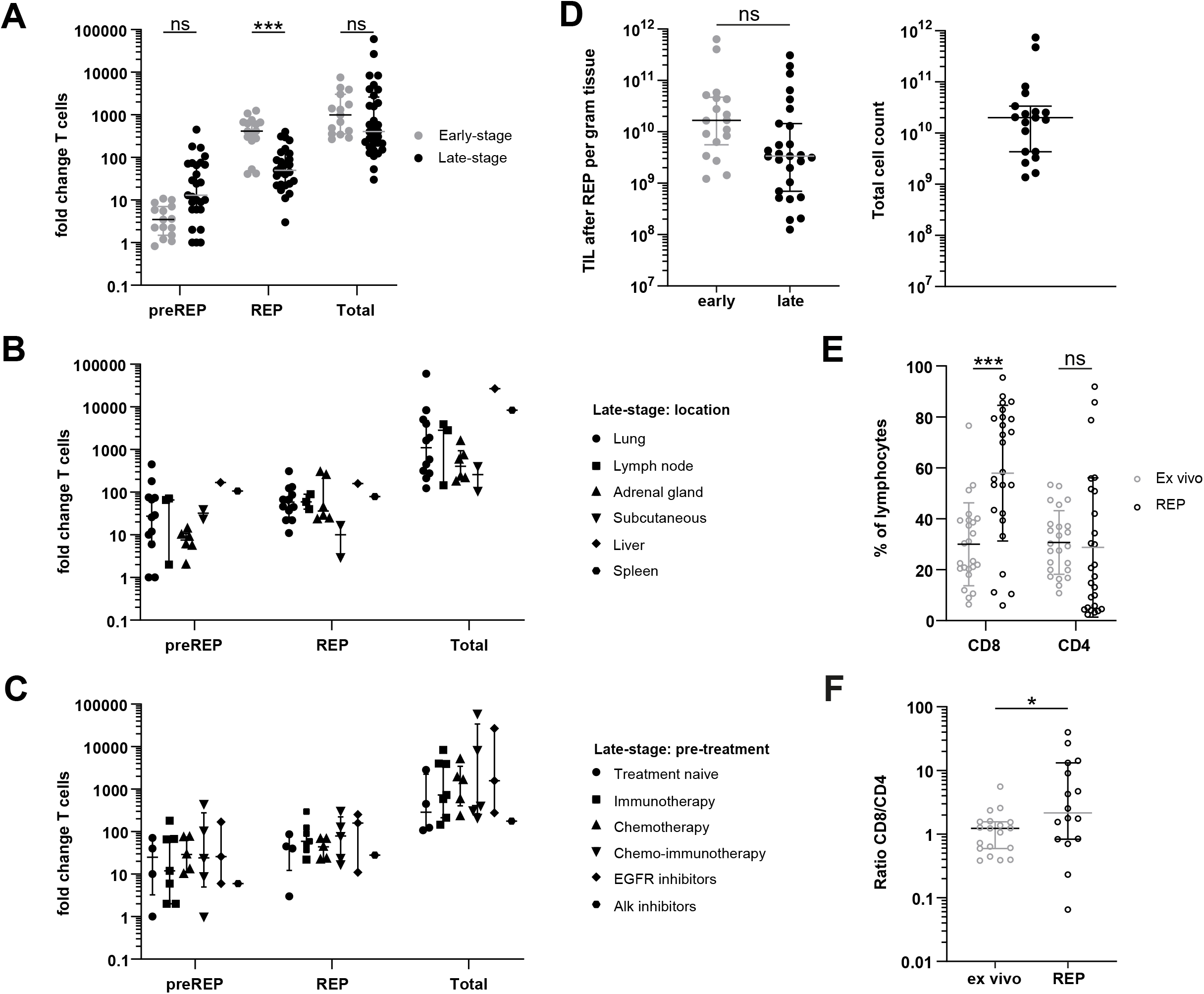
Efficient expansion of late-stage NSCLC TILs. Single cell suspensions from tumor digests were cultured with 6000 IU IL-2 for 10-13 days (preREP), and then for an additional 10-13 days with irradiated feeder cells, 30 ng/ml OKT3 and 3000 IL-2 (REP). (A) Fold change of T cell numbers during preREP (left), REP (middle) and total expansion (right), compared between late-stage (black, n=27) and early-stage (grey, n=15)^14^ NSCLC lesions. The number of life cells was determined with tryphan blue staining. (B,C) preREP (left), REP (middle) and total expansion (right) from (B) different tumor lesion sites and (C) different pre-treatment regimens. (D) Left panel: TIL count per gram tissue for the final TIL product after expansion, compared between late-stage (black, n=27) and early-stage (grey, n=15)^14^ NSCLC lesions. Right panel: total cell count after expansion for late-stage NSCLC lesions, based on the total tumor size and count per gram tissue after expansion. (E) CD8^+^ T cell (left) and conventional CD4^+^ T cell (right) as percentage of lymphocytes, compared between ex vivo (grey, n=27) and REP (black, n=27). (F) ratio of CD8^+^ T cells and conventional CD4^+^ T cells, compared between ex vivo (grey, n=27) and REP (black, n=27). Each dot represents one patient. All graphs show median and 25th and 75th pct interval. Significance was calculated with unpaired student’s t test. * = p<0.05, *** = p<0.001. Not significant (ns): p ≥ 0.05. REP: Rapid Expansion Phase.

We next measured the T cell content of the TIL product at the end of the expansion phase. Similar to our findings in early-stage NSCLC tumor lesions^14^, we observed a loss of Foxp3 expressing T cells after expansion in late-stage TIL cultures (figure S4). The percentage of CD8^+^ T cells increased from 30% ± 16.3% in the tumor digest ex vivo to 58% ± 26.7% REP (p-value 0.006) of the lymphocyte population (figure 5(e); for gating strategy see figure S4). Conversely, the percentage of CD4^+^ T cells maintained constant (figure 5(e)). The ratio of CD8^+^ T cells over CD4^+^ T cells increased from a mean ratio of 1.3 ex vivo to 7.6 after REP (figure 5(f); p-value: 0.04). In conclusion, the expanded TIL product display very high T cell content containing both CD4^+^ TILs and CD8^+^ TILs yet with a higher proportion of CD8^+^ TILs.

### Tumor-responsive T cells are prevalent in TIL products from late-stage metastatic lesions

Lastly, we studied the functionality of the expanded TILs from metastatic NSCLC lesions. To define the overall capacity of TILs products from late-stage NSCLC tumor lesions, we measured their capacity to produce cytokines upon activation for 6 hours with phorbol myristate acetate (PMA) and ionomycin (figure S5). Intriguingly, the capacity of individual TIL products to produce TNF, IFN-γ or IL-2 substantially differed, and some TIL products displayed a clear preference to produce one specific cytokine (figure S5). Nonetheless, each patient produced at least one of the three cytokines to a great extent and most produced two or more (figure S5), revealing their capacity to amply produce cytokines.

To define their tumor reactivity, we next examined the responsiveness of the TIL products to the autologous tumor digest, for which enough material was available from 21 patients. As a read-out, we measured the production of TNF, IFN-γ, IL-2 and CD137 (4-1BB) after 6 hours of co-culture with tumor digest, or with medium alone as control (figure 6(a) and figure S6). To distinguish the expanded TILs from T cell infiltrates of the tumor digest, we pre-stained the expanded TILs with CD4 and CD8 antibodies before the co-culture. The generated TIL products contained not only CD8^+^ T cells, but also CD4^+^ T cells that produced TNF, IFN-γ, IL-2, or that expressed CD137 in response to tumor digest (figure 6(b)). The sum of CD8^+^ T cells that expressed at least one of the cytokines or CD137 (labelled as tumor-responsive) reached 10.0% ± 8.9% in response to tumor digest, compared to 6.6% ± 7.2% positive cells in the medium control (figure 6(c); p-value 0.0007). In CD4^+^ TILs, the increase was even higher and reached 13.0% ± 8.8% of tumor-responsive cells, compared to 4.5% ± 4.3% positive cells in the medium control (figure 6(c); p-value <0.0001). The level of tumor responsive TILs could not be attributed to a specific location or pre-treatment (figure 6(c)).

**Figure 6.**
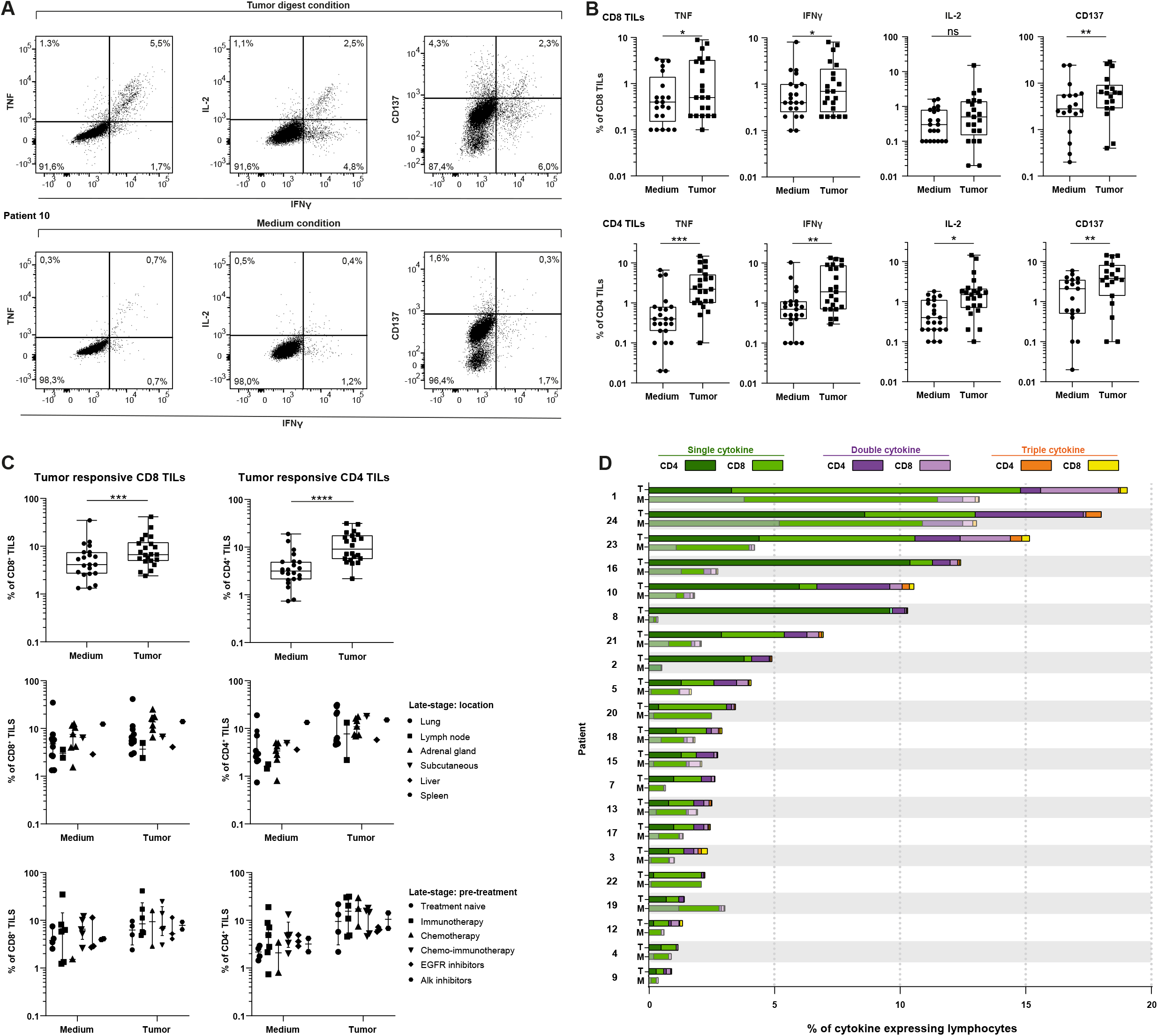
Expanded TILs produce pro-inflammatory cytokines and express CD137 in response to autologous tumor digest. TIL products generated from 22 patients were stained with fluorescently labelled CD4^+^ and CD8^+^ antibodies prior to co-culture with autologous tumor digest or medium for six hours at 37°C. Cytokine production and CD137 expression was defined by flow cytometry. (A) Gating strategy for the TILs exposed to tumor digest (top) or to medium (bottom) (patient 10), based on the expression of IFN-γ (X-axis) and TNF, IL-2 and CD137 (Y-axis). (B) Representative plots showing percentage of positive CD8^+^ (top) and CD4^+^ (bottom) TILs for TNF, IFN-γ, IL-2 and CD137 of tumor-exposed TILs (left)compared to medium control (right) (C) Percentage of tumor responsive CD8^+^ (left panel) and CD4^+^ (right panel) TILs. Tumor responsiveness was defined on the expression of at least one of the markers, i.e. TNF, IFN-γ, IL-2 and CD137 (top), also compared between different tumor lesion sites (middle) and different pre-treatment regimens (bottom). (D) Percentage of cytokine producing TILs that produce one (green), two (purple) or three (yellow) cytokines simultaneously upon exposure to tumor digest (top bar) or to medium (lower bar). Figure A-C: each dot represents one patient. Figure D: each bar represents one patient, patient number corresponds to table 1. Graphs in B and C show median and 25^th^ and 75^th^ pct interval. Significance was calculated with paired student’s t test. * = p<0.05, ** = p<0.01, *** = p<0.001, **** = p<0.0001. Not significant (ns): p ≥ 0.05.

We also defined for each TIL product the number of tumor-reactive CD8^+^ and CD4^+^ T cells that produced more than one cytokine in response to tumor digest. We divided the cells in single cytokine producers, double producers (i.e. that produce two of the three measured cytokines), and triple producers (i.e. that produce all three measured cytokines) (figure 6(d)). Even though the single cytokine producing T cells contribute most to the tumor-responsive TILs, we observed that almost all TIL products also contained double and triple cytokine-producing TILs, and that the level of polyfunctional TILs were in all cases above background (medium control) (figure 6(d)). Notably, we detected these polyfunctional T cells in 16 out of 21 (76%) TIL products, including those that displayed a lower overall percentage of cytokine-producing T cells (figure 6(d)). Thus, even though a large spread of overall tumor responsiveness is found in TIL products from late-stage metastatic tumor lesions, we observed that tumor responsiveness of TIL products as defined by cytokine production and/or CD137 expression is prevalent.

## Discussion

Here, we showed that tumor-reactive TILs can be efficiently isolated and expanded from metastatic NSCLC tumor lesions. This expansion was successful irrespective of the location of the tumor and the pre-treatment regimen the patient received. An important parameter of generating TIL products from metastatic tumor lesions is the presence of T cells. It has been proposed that the origin of tissue from tumors can shape the immune cell composition of metastases^40^. Indeed, the number of CD8^+^ T cell infiltrates in metastatic renal cell carcinomas and colorectal tumors are comparable to those of the primary tumors^41^. With the exception of brain metastasis from NSCLC tumors^42,43^, which were not included in our cohort, the overall lymphocyte infiltration in metastatic NSCLC lesions also closely resembled those of primary NSCLC tumors^14^. This included high T cell infiltrates with a comparable distribution of CD8^+^ T cells, conventional CD4^+^ T cells and regulatory T cells, and similar percentages of B cells, NK cells, and NKT cells. Based on the high T cell infiltration, we thus conclude that all metastatic lesion sites tested could serve as tumor source to expand TILs for therapeutic purposes.

In line with their well-established anti-tumoral cytolytic function^44–47^, the majority of TIL products generated from metastatic NSCLC lesions contained anti-tumoral CD8^+^ T cell responses, as defined by cytokine production and/or CD137 expression after co-culture with autologous tumor digest. Intriguingly, CD4^+^ T cells responded even more robustly to autologous tumor digest, a feature that we did not observe in early-stage NSCLC tumors^14^. Several recent studies identified anti-tumoral cytolytic CD4^+^ T cell responses in solid tumors, including for NSCLC^48–52^. Therefore, we hypothesize that the observed CD4^+^ T cell responses could also substantially contribute to T cell responses against NSCLC tumors. Importantly, 16 out of 21 (76%) TIL products contained polyfunctional T cells, which are considered the most potent anti-tumoral T cells^25,26,53^, further implying that the generated TIL products could achieve potential anti-tumoral responses.

Another important point is whether one could identify tumor lesions with a high likelihood of containing tumor-specific T cells already prior to expansion. We and others previously found a correlation between high PD-1 expression on TILs and anti-tumor responses in TIL products^14,54^. However, circulating anti-PD1/PDL-L1 antibodies due to treatment with nivolumab and/or pembrolizumab (anti-PD-1 immunotherapy) impedes such expression analysis for many of the patients in our cohort. However, other markers may help identify tumor-responsive TILs in such a pre-treated patient cohort. We found that the number of tissue resident T cells as defined by expression of tissue resident markers CD103 and CD69 on CD8^+^ T cells and by CD69 expression on CD4^+^ T cells was increased in late-stage tumor lesions. These tissue-resident TILs showed increased expression of CD39 and were enriched for CD137 expression, which is indicative for TCR engagement and thus potentially for tumor-specific TILs^37,39,55^. CD137 positive T cells were shown to express the highest levels of IFN-γ, TNF, IL-2^39,56,57^, which further points to their capacity to respond to tumor cells. It is therefore tempting to speculate that the combination of these markers, i.e. CD103, CD69, CD39 and CD137 could identify, and potentially even enrich for tumor-specific TILs.

TIL therapy has demonstrated its potential in clinical trials for late-stage melanoma patients^20,22,58^, bladder cancer^59^, breast cancer^60^, ovarian cancer^61^, head and neck cancer^62^, and recently also for NSCLC^27^. In this phase I clinical trial for anti-PD-1 refractory NSCLC patients, 11 out of 16 patients (76%) showed an objective response, and two a complete response, which highlights the potential of TIL therapy for NSCLC patients^27^. Importantly, as for melanoma alike^63^, this study also revealed that TIL therapy can be beneficial for patients that failed immunotherapy. Our results indicate that neither a specific pre-treatment regimen, or the tested locations of metastatic lesions should be a reason to exclude a patient from receiving TIL therapy. However, because the efficacy of TIL therapy is reduced when used as second or third treatment regimen^64^, it may be key for effective treatment to identify which therapy is most suitable for a patient, and to consider TIL therapy as first line treatment.

In conclusion, we showed here that the generation of tumor-reactive TIL products from late-stage NSCLC tumor lesions is feasible, irrespective of pre-treatment regimen or tumor origin. The obtained TIL products reach sufficient cell numbers for clinical application, and most products contain T cells with polyfunctional anti-tumor effector functions. We are therefore confident that TIL therapy will generate promising results for metastatic NSCLC patients in upcoming clinical trials.

## Authors contribution

SMC, RdG, KM, KJH, EFS, JBAGH and MCW designed the study; SMC, RdG and AG performed experiments and analysed data, KJH and AAFAV coordinated sample collection and performed pulmonary surgery, KM collected and analysed samples. SMC and MCW wrote the manuscript. MCW supervised the study. All authors read and approved the final manuscript.

## Acknowledgments

We thank the Flow cytometry facility from Sanquin Research, and the medical assistance staff from the NKI - AvL for technical help.

## Ethics approval and consent to participate

The study was performed according to the Declaration of Helsinki (seventh revision, 2013), and executed with consent of the Institutional Review Board of the Netherlands Cancer Institute-Antoni van Leeuwenhoek Hospital (NKI-AvL), Amsterdam, the Netherlands (CFMPB317).

## SUPPLEMENTAL FIGURE LEGENDS

**Supplemental Figure 1. Ex vivo gating strategy to study lymphocyte infiltrate from late-stage metastatic tumor lesions.** Top left panel: Dead-cell exclusion with Near-IR viability dye. Top right panel: lymphocyte inclusion based on FSC-A and SSC-A. Bottom left panels: single cell inclusion based on SSC-H, SSC-W and FSC-H, FSC-W. Bottom right panel: CD3^+^ T cell inclusion.

**Supplemental Figure 2. Effective isolation of lymphocytes from late-stage NSCLC lesions.** (A) Flow cytometric determination of lymphocyte infiltrates in single cell suspensions per gram tissue from late-stage metastatic NSCLC lesions (black, n=25) compared to early-stage NSCLC lesions (grey, n=17)^14^. (B) Lymphocyte infiltration in late-stage NSCLC tumor lesions at different metastatic lesion sites. (C) Comparison of different pre-treatment regimens the patients received prior to metastatic tumor excision. Each dot represents one patient. All graphs show median, 25th and 75th pct interval. Significance was calculated with unpaired student’s t test. Not significant (ns): p ≥ 0.05.

**Supplemental Figure 3. CD39 and CD69 expression in CD8^+^ T cells ex vivo.** (A) Gating strategy based on CD39 and CD69 expression (patient 13). (B) CD39 and CD69 expression on CD8^+^ T cells. (C) CD137 (4-IBB) expression, calculated as percentage of the subpopulation labelled on the X-axis in CD8^+^ T cells. Each dot represents one patient. Graph shows median, 25th and 75th pct interval. Significance was calculated with 2-way ANOVA test (Tukey’s Multiple Comparison). * = p<0.05, ** = p<0.01, *** = p<0.001, **** = p<0.0001. If no indication, p ≥ 0.05.

**Supplemental Figure 4. REP gating strategy**. Expanded TILs were collected, and phenotypic analysis was performed by flow cytometry. Top row, left panel: dead-cell exclusion with Near-IR viability dye. Middle panel: lymphocyte inclusion, based on FSC-A and SSC-A. Right panel, and Bottom row, left panel: single cell inclusion based on SSC-H, SSC-W and FSC-H, FSC-W. Middle panel: CD4^+^ TIL and CD8^+^ TIL selection. Bottom right panel: Treg selection based on Foxp3 and CD25.

**Supplemental Figure 5. Expanded TILs are functional and capable of producing cytokines after stimulation with PMA and ionomycin.** For both CD8^+^ (left) and CD4^+^ (right) TILs, cytokine production and CD137 expression was determined with flow cytometry after PMA and Ionomycin stimulation for six hours at 37°C. Each dot represents one patient. Median and 25th and 75th pct interval is shown. The dot plots show flow cytometry data from patient 12, and 18.

**Supplemental Figure 6. Gating strategy for tumor reactivity assay for two different patients.** TILs from different patients were pre-stained with fluorescently labelled antibodies for CD4^+^ and CD8^+^ and co-cultured with autologous tumor digest or medium as a negative control for six hours at 37°C. TILs were stained with fluorescently labelled antibodies to determine the tumor reactivity with flow cytometry. Gating strategy for the TILs co-cultured with autologous tumor digest (top) or medium (bottom) for an additional two patients; (A) patient 11 and (B) patient 1.

