## Supplementary figures and images for "Effective generation of tumor-infiltrating lymphocyte products from metastatic non-small-cell lung cancer (NSCLC) lesions irrespective of location and previous treatments"

### Supplemental figure 1

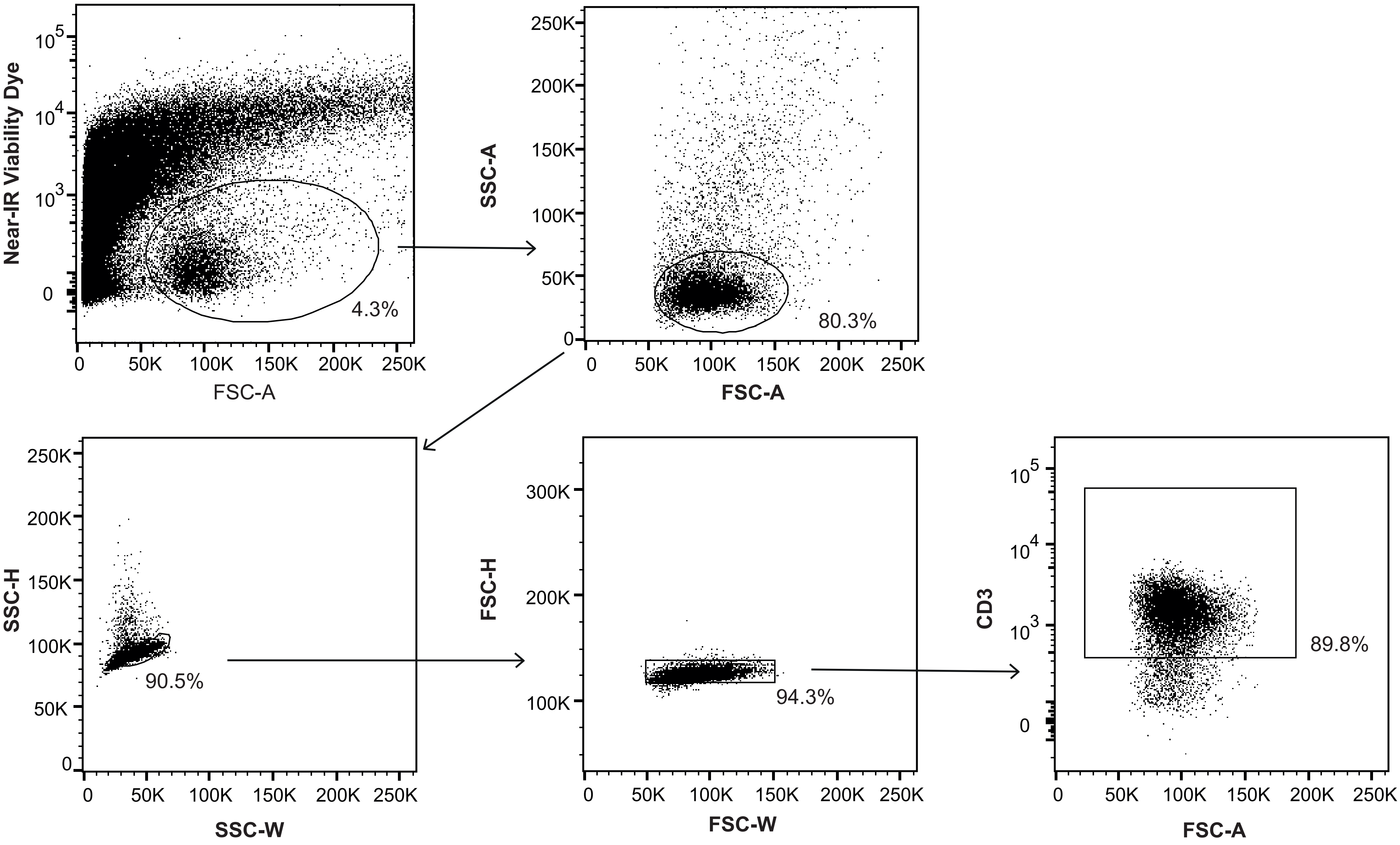

### Supplemental figure 2

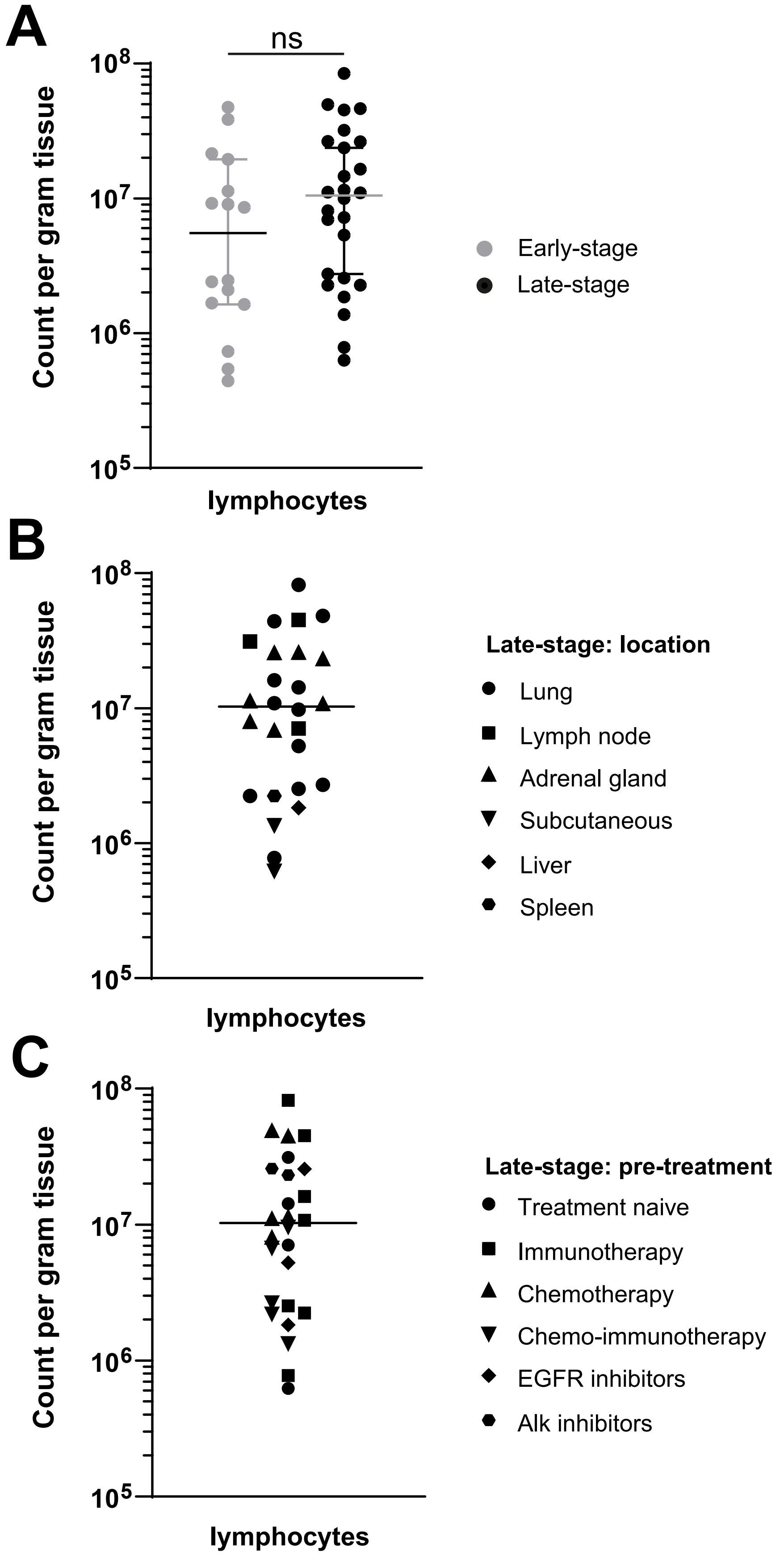

### Supplemental figure 3

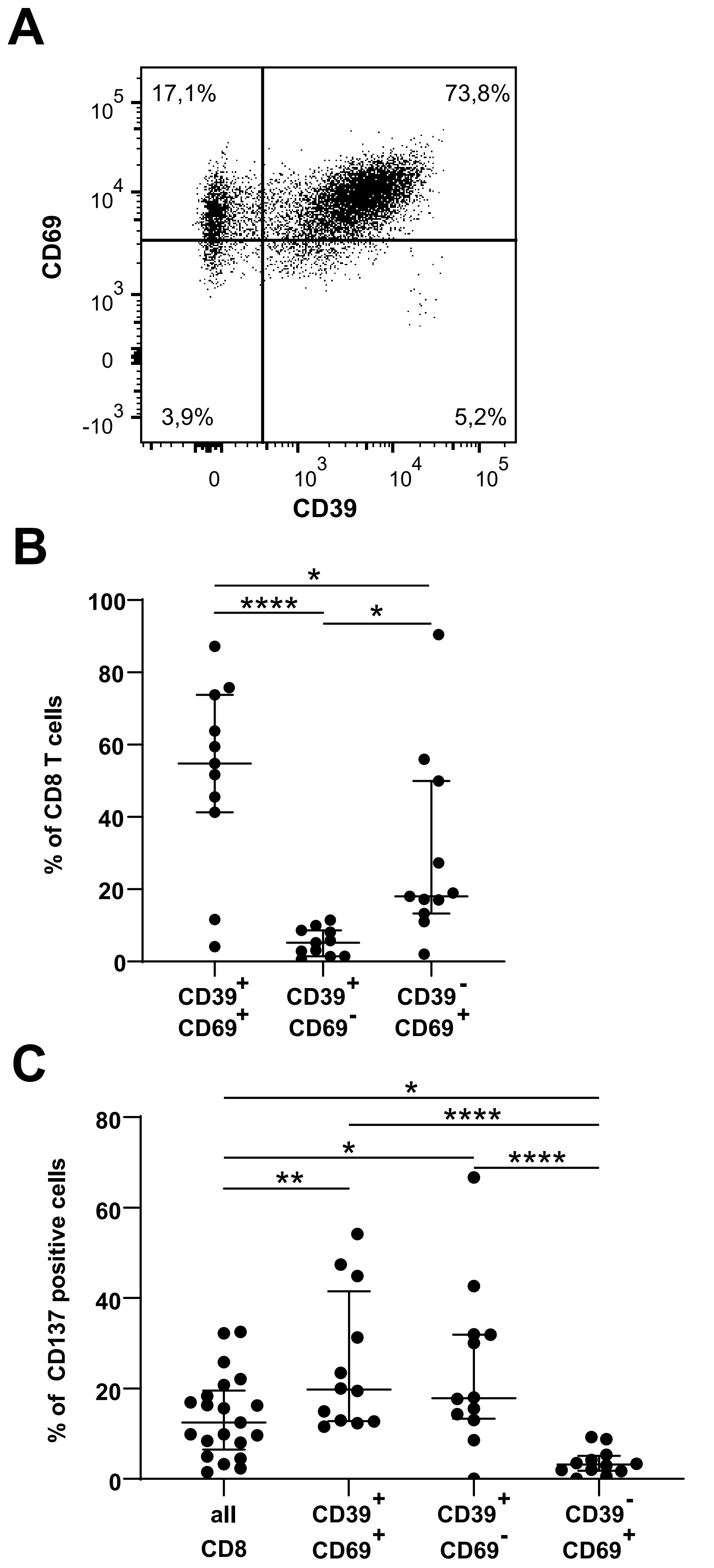

### Supplemental figure 4

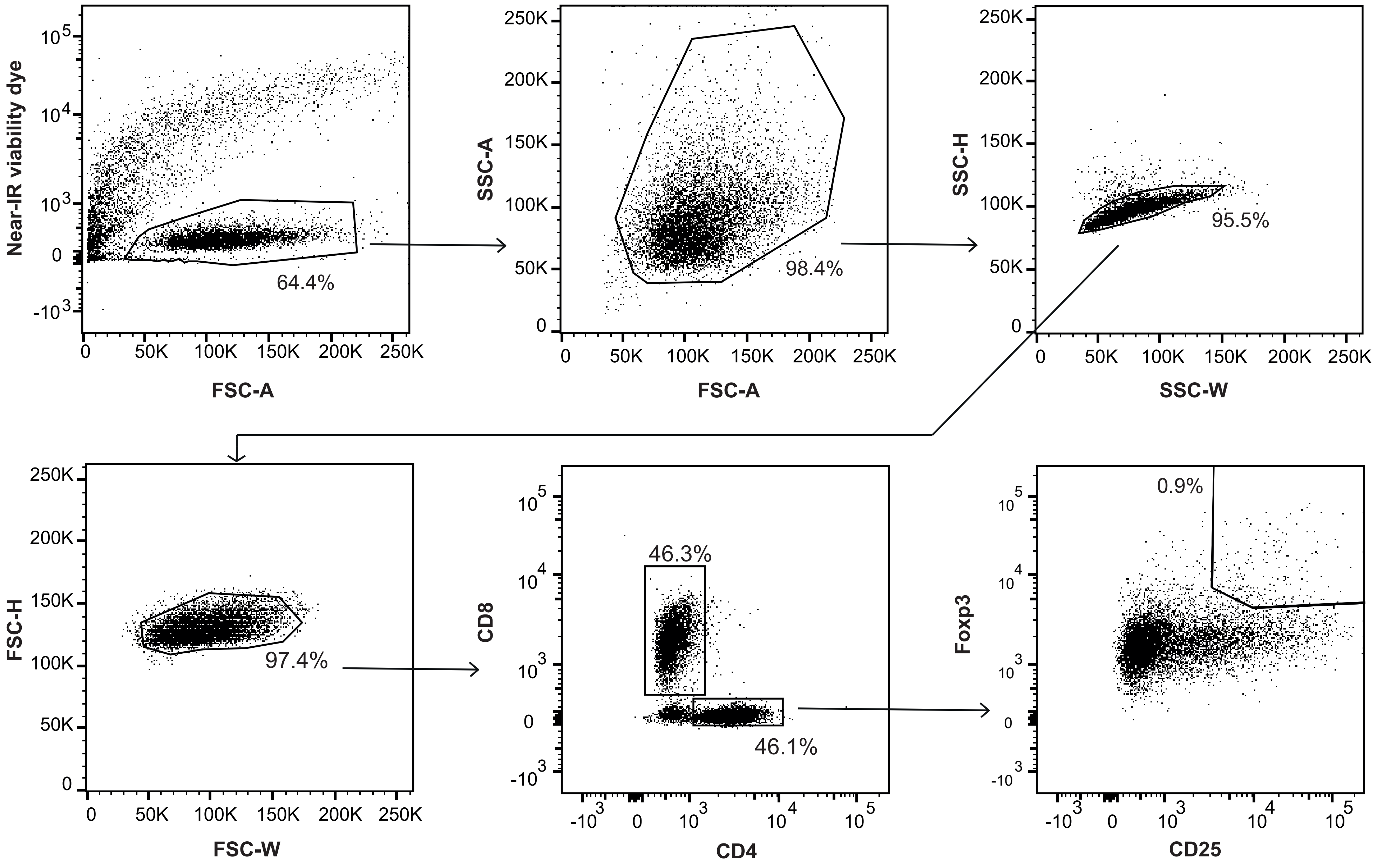

### Supplemental figure 5

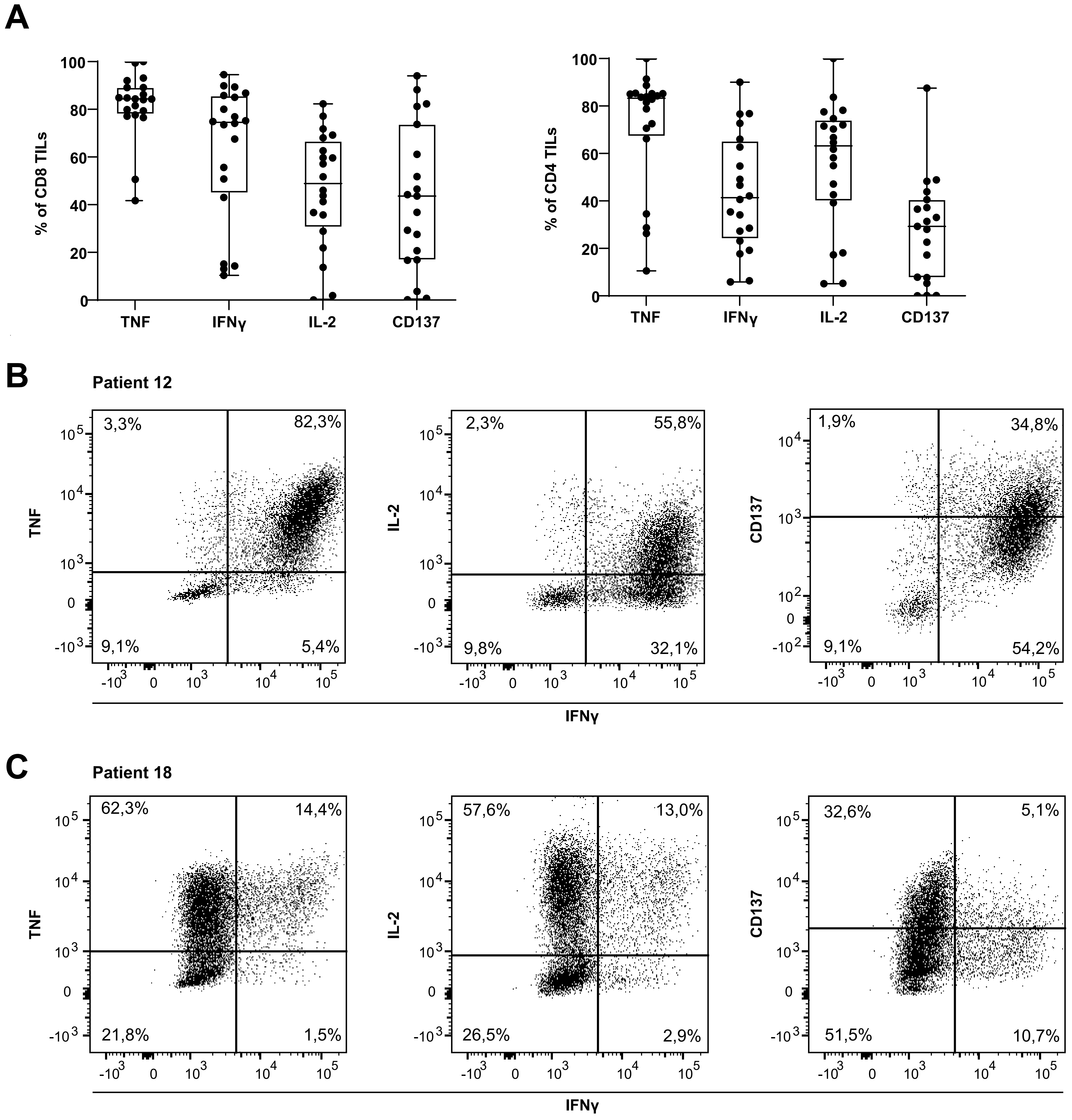

### Supplemental figure 6

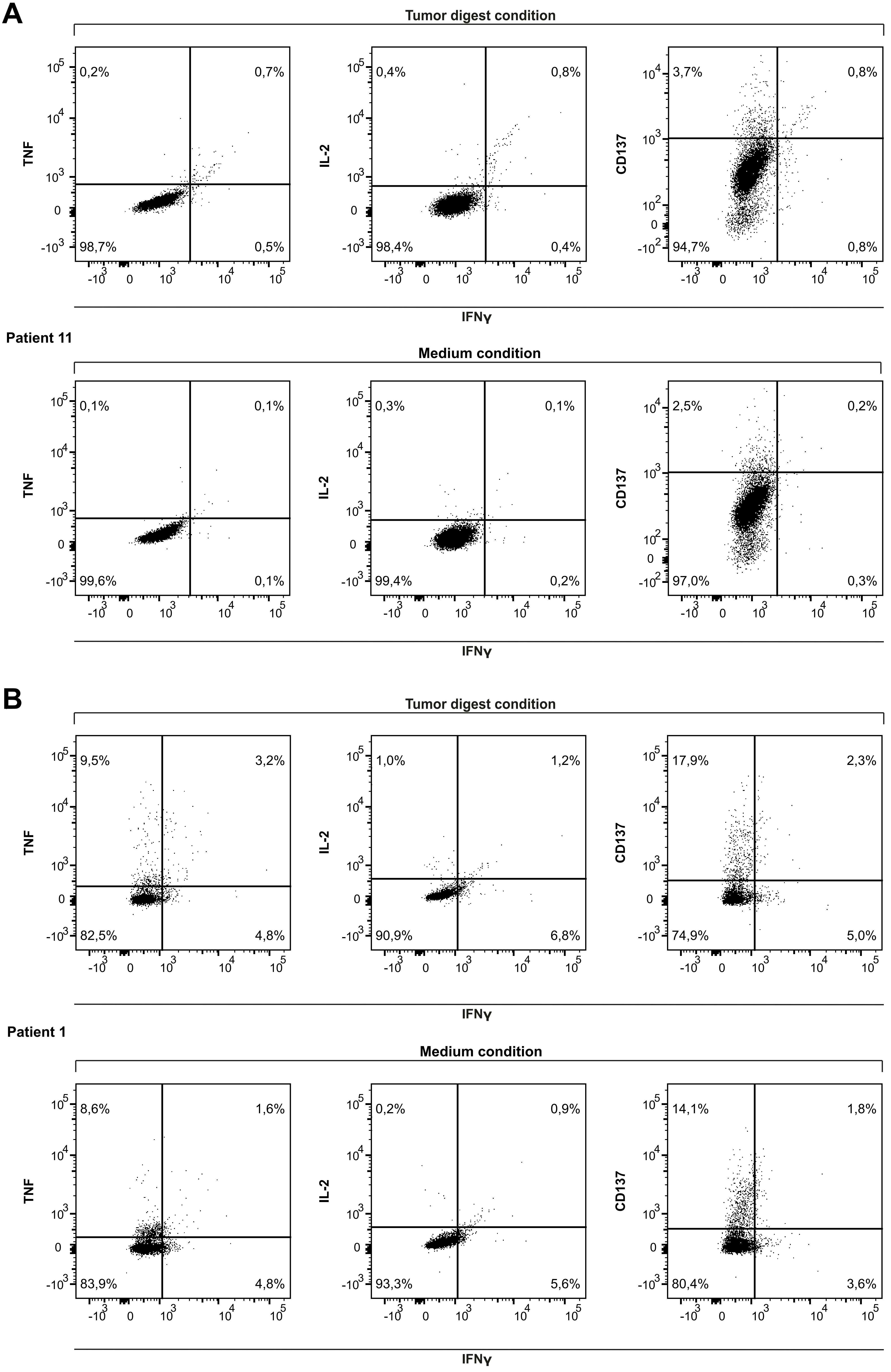
